# Analysis of the economic viability and environmental impacts of a conceptual process for the recovery of lactic acid from spent media in cultivated meat production

**DOI:** 10.1101/2025.04.30.651479

**Authors:** Josh Wimble, Reina Ashizawa, Elliot W. Swartz

## Abstract

The scaled production of cultivated meat implies a future where large amounts of liquid waste in the form of spent media will be co-produced. Recycling of spent media, specifically certain abundant metabolites such as lactic acid, offers an opportunity for valorization and to offset the carbon footprint of cultivated meat production, however, the feasibility of recovering lactic acid from spent media has yet to be examined in detail. In this study, we developed a conceptual design of a five-step lactic acid recovery process integrated into a previously modeled cultivated meat facility. We examined the corresponding cost and environmental impacts of recovering an 88% aqueous, polymer-grade lactic acid solution and compared these footprints to data from commercial lactic acid fermentation processes. At an anticipated lactic acid concentration in spent media of 3 g/L, we found that the net cost of recovery would be $0.71 per kg of 88% lactic acid, with a 7.5 year simple payback period. Sales of this co-product could offset $0.06/kg of the cost associated with the production of cultivated meat. Depending on allocation scenarios, the environmental impact of the recovery process had a-1.0 to +0.2 kg CO_2_ eq effect on the overall carbon footprint and a-22 to +3 MJ effect on cumulative energy demand per kg of cultivated meat production. These results suggest that the recovery of lactic acid may be an economically viable and environmentally beneficial practice if implemented in future production facilities. This study provides crucial guidance for lactic acid valorization and other media recycling strategies for cultivated meat that can be applied to broader animal cell biomanufacturing industries.

## Introduction

Cultivated meat is a nascent technology that involves cultivating animal cells under controlled conditions to create meat and seafood products without the need to raise animals. Cell cultivation occurs in a medium composed of nutrients such as amino acids, glucose, vitamins, and inorganic salts, often supplemented with added factors such as antioxidants, lipids, and growth factors.

The cell culture medium is the largest cost and environmental impact driver of production ^1,2^. The degree to which media drives cost and environmental impacts depends on how the media ingredients are sourced and how efficiently they are used. Media use efficiency is influenced by cellular metabolism (the feed conversion ratio), process design, recycling ^3^, and bioreactor operation.

Glucose is the most abundant nutrient in the media by weight. During rapid proliferation in bioreactors, cells typically undergo anaerobic respiration known as the Warburg Effect, resulting in the conversion of glucose into lactic acid. Models of different scenarios for media use efficiency for cultivated meat production suggest that between 256 and 401 kg of lactic acid could be produced for every ton of cultivated meat ^2^. Given the cost-sensitivity and motivation of sustainability in the cultivated meat sector, this amount of lactic acid makes it an attractive target for valorization as a co-product, displacing virgin lactic acid production for use in downstream industries such as bioplastics and food. One study estimated that if lactic acid was captured as a co-product and allocated based on mass, it could offset the carbon footprint of cultivated meat production by 63 to 85% ^4^. However, the techno-economics and energy requirements of such recovery and how they compare to commercial lactic acid fermentation have yet to be analyzed.

Commercial lactic acid fermentation is a multi-billion dollar market that is expected to produce nearly two million tons of product annually by 2025 ^5^, with commodity prices varying by grade and concentration, generally between $1300-2300 per metric ton ^6^. Accordingly, many well-studied technologies are available for the recovery of lactic acid from aqueous fermentation streams, including extraction (with and without reactive extractants), adsorption, chemical methods such as esterification and reactive distillation, and membrane separations, including reverse osmosis and nanofiltration ^7^. Improvements for each of these techniques continue to evolve, with new approaches in development such as resin wafer electrodialysis^8^ and novel resins using RNA to increase the specificity of lactic acid recovery ^9^.

The recovery of lactic acid from spent media differs in two key ways from lactic acid fermentation. First, the concentration of lactic acid is expected to be between 1 and 3 g/L ^10^, which is significantly less than commercial fermentation concentrations that frequently exceed 100 g/L ^11^. 3 g/L (33.3 mM) is the concentration at which lactic acid poses growth inhibition for many animal cell lines, and cultivated meat bioprocesses are likely to hold lactate concentrations near this boundary ^12^. Second, spent media from cultivated meat production is a complex solution with a broader range of metabolites and ingredients than microbial fermentation media. These two factors imply that additional processing steps may be required to yield purified lactic acid compared to existing commercial fermentation processes. Notably, although the complex makeup of spent media presents challenges, one advantage is that animal cells naturally produce L-lactic acid, which is the preferred isomer for poly-lactic acid (PLA) synthesis.

In this study, we demonstrate that dilute lactic acid contained within spent media could be cost-effectively purified, enabling a new revenue stream for manufacturers while avoiding the upstream footprint associated with microbial fermentation to produce virgin lactic acid. We developed and modeled a theoretical five-step recovery process using existing separation and purification technologies and calculated the estimated costs, carbon footprint, energy demand, and water use for this process when integrated into a previously modeled hypothetical facility producing 10,000 tons of cultivated meat annually ^2^. Finally, we compared these values to bulk pricing and environmental impact data for commercial lactic acid production to determine its potential viability in the real world.

## Methods

### 1. Composition of spent media and purification considerations

Spent media contains nutrients, metabolites, growth factors, and other biomolecules (e.g., exosomes, nucleotides) that were either not consumed or were secreted by the cells during the duration of the cell culture. The most abundant molecules in spent media are typically glucose and lactic acid, which can be measured at low g/L concentrations. Amino acids and vitamins are also relatively abundant, measured at mg/L concentrations, while growth factors can be measured in pg/L to ng/L ^10^. Salts and minerals are typically not consumed at high rates and remain in the spent media at concentrations up to mg/L ^10,13^. **Table 1** summarizes key classes of molecules in spent media and their respective attributes relevant to industrial separation, with additional considerations described in the **Supplementary Information.** Taken together, a process that aims to recover and purify lactic acid from spent media will require multiple unit operations to separate impurities from other spent media components and avoid downstream complications.

**Table 1.**
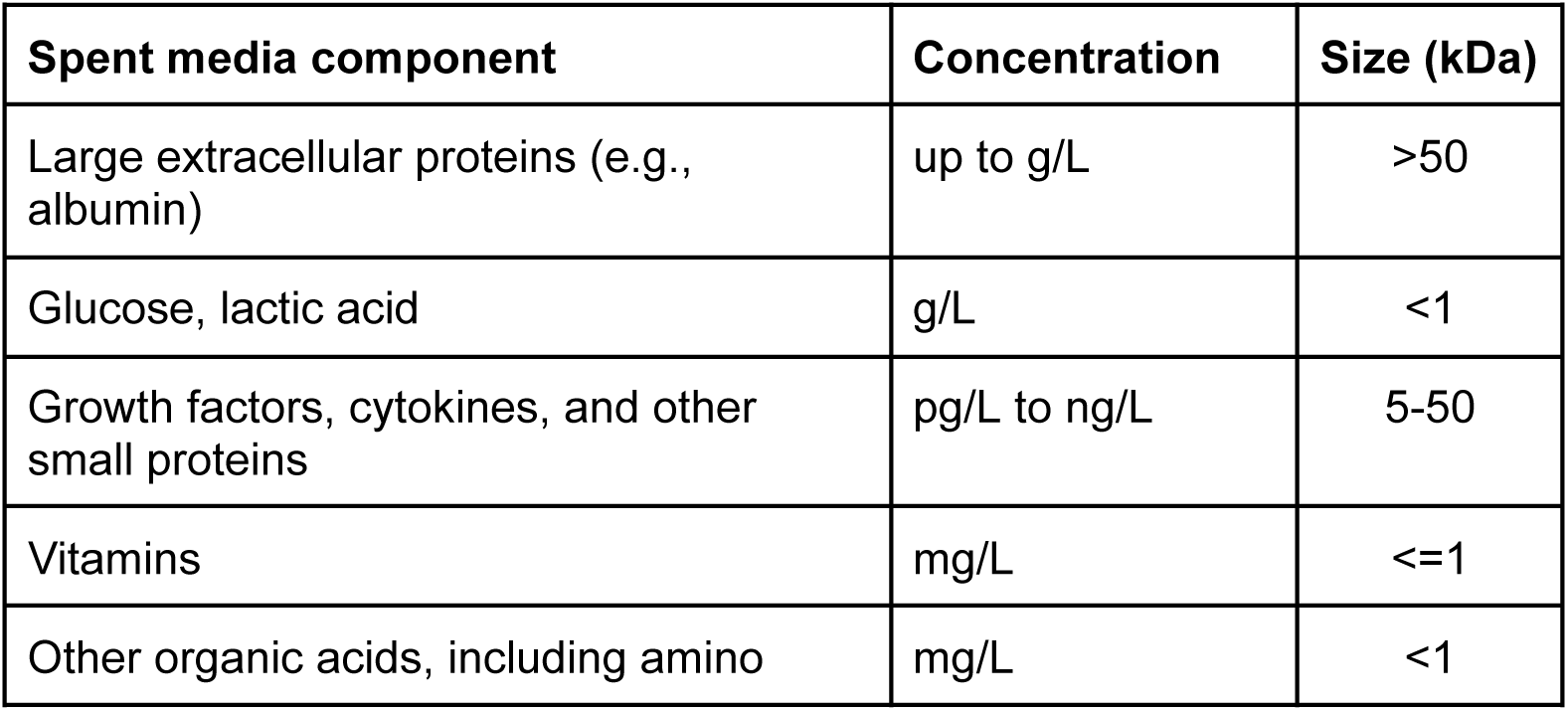
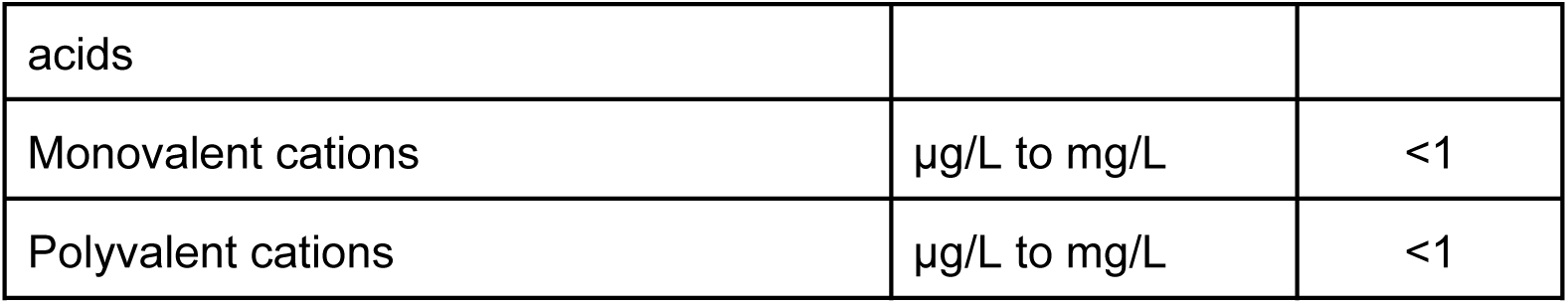
Expected concentrations and sizes of key classes of molecules found in spent media.

### 2. Comparison of existing process technologies

Because lactic acid recovery from microbial fermentation has been the subject of extensive research, a wealth of data on the recovery of lactic acid from fermentation broth at moderate concentrations ranging from approximately 10-200 g/L is available ^14^. Suitable solutions for lactic acid recovery from spent media were investigated, culminating in a proposed recovery process that could be implemented in a hypothetical cultivated meat manufacturing facility.

### 2.1 Extraction

Extractive techniques have been studied extensively for the continuous separation of lactic acid from fermentation broth to improve lactic acid yields by limiting the inhibitory effects of lactic acid and lactate on cells ^15,16^. Extractive techniques use an immiscible solvent and, optionally, a chemical extractant to extract lactic acid or lactate from the aqueous phase, followed by lactic acid recovery, commonly by distillation or back extraction into a second aqueous phase. While continuous extraction has shown promise for increased lactic acid productivities and yields, its practical utility to cultivated meat production is limited by low partition coefficients in common immiscible solvents at physiological pH due to lactic acid’s hydrophilic nature. Further, even ppm levels of common extractants such as tertiary amines may be detrimental to cell growth and viability ^17^, and residual extractants may pose food safety issues.

Indirect or offline lactic acid extraction provides some advantages for cultivated meat processes because the offline treatment of spent media would not require direct contact between extractants and cells, and a wider selection of less toxic solvents and extractants would be available for consideration. Further, spent media may be pH adjusted to less than the pKa of lactic acid, which can significantly improve solvent partitioning without concern for impacts on cells. However, this approach does not address the other challenges with extractant and solvent removal.

While solvent extraction is expected to provide good separation of lactic acid from residual proteins and salts, poor separation of glucose and other macromolecules in spent media would also be expected, resulting in these neutral molecules being coextracted with lactic acid. Given these challenges, extraction was not prioritized for the model recovery process.

### 2.2 Membrane separation

Membrane separations are used in many industrial processes such as desalination and sugar concentration. They are attractive for lactic acid concentration and recovery because they are generally more energy-efficient than thermal separation processes ^18^ and their capital and operating costs are well-characterized ^19^.

Many different membrane technologies are available for the separation and concentration of target molecules, ranging from reverse osmosis (RO) to microfiltration (MF) and more recently electrodeionization, depending on the properties of the solute of interest. Other research has proposed purely membrane-based systems for the recovery of lactic acid from complex feedstreams, including spent media ^4^. The separation mechanisms of electrolytes and neutral solutes are well understood for membrane filtrations, and nanofiltration (NF) provides promising separation characteristics for spent media components such as glucose and monovalent ions from lactic acid through a combination of size sieving and Donnan exclusion effects ^20^.

In addition to the use of NF for lactic acid recovery, membrane filtration technologies provide promising routes to separate large species such as cell debris, proteins, and other macromolecules, with varying rejection rates depending on the membrane properties and application. Given this context, MF and NF membrane separation steps were selected for the model lactic acid recovery process.

### 2.3 Adsorption

Adsorption of lactic acid from fermentation broths has been explored extensively as an alternative to traditional lactic acid recovery techniques used in microbial fermentation ^3,7^. Because of the moderate pKa of lactic acid, two separation approaches are possible. In the first approach, spent media is processed above the pKa of lactic acid without pH adjustment, such that the predominant species is the lactate anion. In this operating mode, Amberlite® resins have been considered as model absorbents with well-understood capacities and recoveries, and have been used to remove metabolites for media recycling in cultivated meat company patents ^21^.

In the second approach, the spent media is pH-adjusted to below the pKa, and separation is based on physical adsorption by the absorbent. This approach has been explored extensively, and both operating modes have been demonstrated for the recovery of lactic acid, with the preferred route depending on the choice of absorbent ^22^ ^23^.

Ion exchange (IEX) resins and other adsorbents also have a long history of use for contaminant removal and polishing steps. For example, activated carbon is used in a variety of industrial processes, including decolorization in sugar refining and plastics recycling, and has been noted by other researchers for its potential applicability in media recycling ^3^. The use of activated carbon has also been investigated for the separation of sugars from acidic solutions produced during the hydrolysis of starchy biomass as feedstock for bio-ethanol and lactic acid production, which requires the removal of sulfuric acid before use ^24^. Accordingly, activated carbon and IEX were selected for the model lactic acid recovery process.

### 2.4 Direct evaporation

While final lactic acid specifications vary by application, concentration by distillation or evaporation is usually required to produce concentrated solutions for use or further processing. As is the case with many industrial processes, this final processing step is expected to be the most energy-intensive unit operation in the recovery of lactic acid from spent media, and variants of this technology to reduce energy demand are well developed.

For example, vacuum-assisted evaporation used in multiple effect evaporators (MEE) is a mature technology for water removal with the ability to drastically reduce energy consumption through the use of reduced pressure and latent heat recovery from distillate streams. Vacuum evaporation has been demonstrated in recovery processes for high-purity lactic acid from fermentation broths ^20^ and the use of three to seven effect evaporators is common in the pulp and paper industries for the concentration of weak black liquor, where steam economies (kg of water evaporated per kg of live steam supplied) above 6 have been shown ^25^.

Similar approaches are expected to provide important energy reduction for the recovery of lactic acid from spent media, though special construction materials must be considered due to the corrosivity of high-concentration lactic acid. It’s feasible that a well-designed cultivated meat plant with heat integration could further reduce the energy usage associated with evaporation, however, heating requirements for large-scale cultivated meat production are predominantly for pre-heating media, so only a small portion of waste heat is directly usable within cultivated meat production ^2^.

One additional benefit of evaporative techniques for concentration is the possibility of recovering water for reuse within the facility, as the distillate streams from evaporation are expected to be free of major media components and appropriate for reuse as water for CIP or for media makeup following polishing to remove residual lactic acid.

## 3. Conceptual process for lactic acid recovery and purification

High-purity lactic acid is required as the raw material for downstream polymerization because of the sensitivity of the PLA production process to impurities. For this study, we used 88% lactic acid as the target co-product, as this concentration is suitable for a range of downstream applications, including food, cosmetics, and PLA production. To achieve this concentration from dilute lactic acid present in spent media, the process first requires the separation and recovery of lactic acid from four distinct categories of species in spent media prior to purification via evaporation: 1) small and large proteins and cell debris, 2) small, generally polar organics such as glucose, vitamins, and other organic acids, 3) mineral salts added to maintain osmotic balance and to support enzyme function, and 4) other organic acids.

**Table 2** provides a high-level overview of the separation capabilities of the techniques previously discussed, based on the expected partitioning of lactic acid. From these separation capabilities and previously described considerations, a five-step recovery process was derived to address each category of molecules for separation, followed by the final concentration of lactic acid (**Figure 1**).

**Figure 1.**
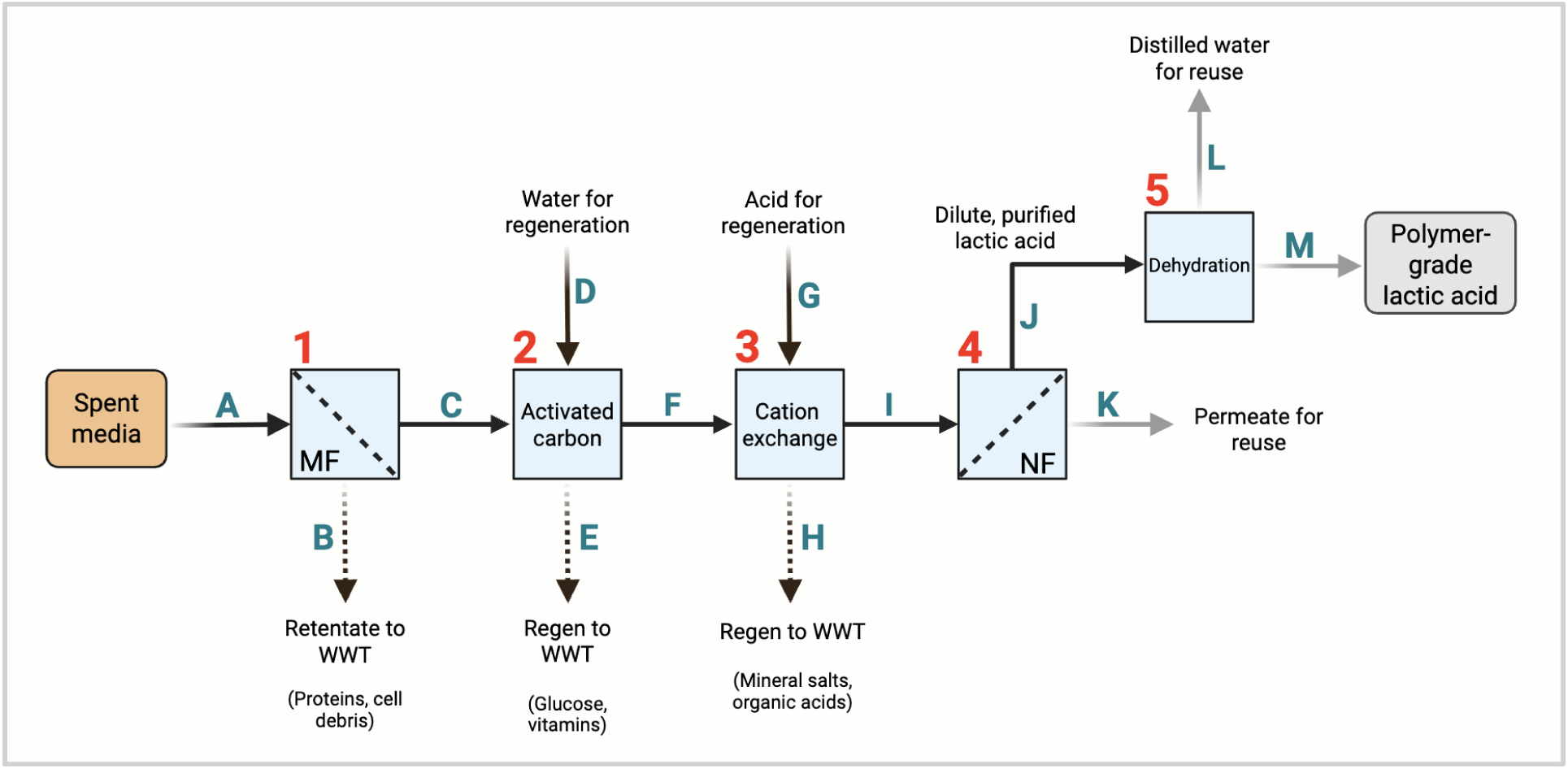
A proposed five-step process for the separation and purification of lactic acid from spent media. Numbers denote individual unit operations, and letters denote material flows, which are described in the text. WWT = wastewater treatment; MF = microfiltration; NF = nanofiltration.

**Table 2.** Overview of the separation capabilities of existing process technologies.

|  | Small and large proteins | Polar organics | Salts | Other organic acids |
| --- | --- | --- | --- | --- |
| **Extraction** | High | Low | High | Low |
| **Membrane techniques** | High | Low | Low | Low |
| **Adsorption (high pH)** | High | High | Low | Low |
| **Adsorption (low pH)** | High | Low | High | Medium |
| **Evaporation** | None | None | None | None |

## 4. Material balance and model construction

The recovery process was assumed to be integrated into the cultivated meat manufacturing process (**Figure S1**) from a previous study where environmental impacts were analyzed in a hypothetical facility producing 10,000 metric tons annually (10k MTA)^2^. The baseline scenario for media use was selected, corresponding to 144,382 L per 3,080 kg batch (46.9 L of media per kg of cultivated meat). These values were used to estimate a spent media production rate of 58,597 L per hour, assuming 8,000 facility operating hours per year.

It was determined that the baseline scenario in Sinke et al results in lactic acid concentrations of 6.8 g/L, which may pose viability issues without additional cell engineering or process control strategies. Accordingly, the baseline lactic acid concentration in this study was set at 3 g/L in the spent media stream entering the recovery process, which is around the setpoint that most cultivated meat bioprocesses are expected to be held at or below ^12^. Ammonia concentrations in spent media were calculated from the same dataset. Since glucose in spent media is typically not entirely depleted, concentrations from published studies using different cell lines were averaged to arrive at a value of 1.8 g/L that was included in the model.

Using these values, the material balance and model were established (**Figure 2**). Equipment size and utility usage were derived based on typical operating conditions and required flow rates for each process step. Capital and operating costs were then calculated and used to derive the direct cost of goods sold (COGS), while materials and utilities were used to establish an inventory for life cycle assessment (LCA). The full model and calculations can be found in the **Supplementary Spreadsheet**.

**Figure 2.**
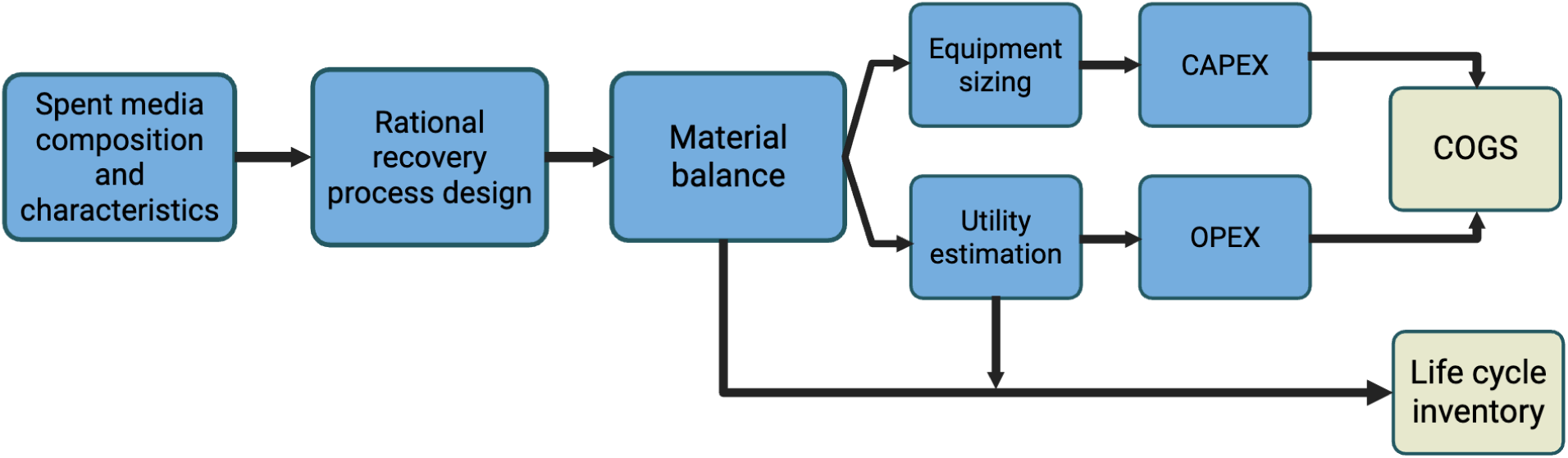
Overall model construction showing how spent media composition is used to inform cost and life cycle inventory estimates for the proposed recovery process. CAPEX = capital expenses; OPEX = operational expenses.

## 5. Recovery process overview and assumptions

In the proposed recovery process outlined in **Figure 1**, spent media (flow A) enters the first step, where crossflow membrane filtration is used to remove cell debris and extracellular proteins from the spent media. The permeate containing glucose, vitamins, mineral salts, and lactate is collected in a receiving tank for further processing (flow C). A 15x concentration factor was assumed such that the total retentate flow to wastewater treatment (flow B) was approximately 3,900L/hr.

In the second step, the clarified permeate (flow C) is processed through duplex activated carbon beds to remove organics such as glucose. The bed size was calculated based on the adsorption characteristics of glucose as measured by Park et al., and the duplex arrangement allows for the loading of one bed while the other is regenerated ^24^. The modeled regeneration step assumed five bed volumes of recycled water (flow D) from the NF and evaporation steps later in the process, which is then sent to wastewater treatment (flow E). Eluate from step two (flow F) is collected in the feed tank prior to step three.

In the third step, IEX is performed using a similar equipment arrangement as in step two. The IEX bed was sized assuming a resin capacity of 2eq/L, and the total media cation equivalents were calculated based on typical DMEM/F12 formulations. IEX regeneration assumed three bed volumes of 1M H_2_SO_4_ (flow G) to recover the acid form of the IEX resin after treatment, which is then sent to wastewater (flow H). Purified lactic acid is then collected in the NF/RO system feed tank.

In the fourth step, a three-stage NF/RO setup was modeled assuming 95% lactic acid retention. This configuration uses sequential membranes to allow progressive concentration of lactic acid by dewatering the feed solution. Membrane areas and flow rates in each stage were calculated based on typical permeate fluxes. Membrane replacement costs and LCA impacts were estimated from available data, assuming a five-year membrane lifetime. In this unit operation, lactic acid concentration increases to 77 g/L (flow J) with an overall lactic acid yield of 86%. The NF retentate is collected in a receiving tank ahead of evaporation. The permeate water (flow K) then becomes available for reuse within the facility.

In the fifth step, a five-effect evaporator was modeled to concentrate the resulting purified lactic acid stream to 88% polymer-grade lactic acid. 88% was taken as the target concentration because it’s commonly used in food additive and pharmaceutical applications. While higher concentrations are possible, further dehydration can lead to the formation of oligomers, which present challenges in producing higher purity lactic acid. Typical heat transfer coefficients were used to estimate the required evaporator area, and an assumed steam economy of 4.5 was used to calculate steam use for water removal. Heat integration within the plant was not considered, and only the steam required for evaporation was included in the utility usage to approximate a system in which waste-sensible heat from the MEE is used to preheat the feed stream. The resulting 88% lactic acid (flow M) is collected in a product storage tank, and capital was not included for truck loading or other final disposition. Recovered distilled water (flow L) is considered to be available for reuse within the lactic acid separation process or to displace RO water usage in the integrated facility for uses such as media makeup.

Regarding water use, the baseline scenario from Sinke et al assumed that 75% of water was reused ^2^. In this study, water and wastewater are expressed in terms of net usage to highlight the change in use with an integrated recovery process versus the reference case.

## 6. Costs

Equipment costs were estimated based on available correlations and were adjusted to 2023 dollars using Chemical Engineering Plant Cost Index (CEPCI) data available from ^26^. Typical installation factors were included for process equipment available from Couper et al ^27^, but other direct and indirect costs associated with the installation, including buildings, engineering, contingency, and startup and commissioning, were not included. While these costs are important in calculating the overall fixed capital investment of the proposed process, they are difficult to estimate for a hypothetical 10k MTA cultivated meat facility due to uncertainty around greenfield and brownfield decisions and overall facility and process integration. Major equipment items were included in the capital cost estimates, but incremental costs associated with utility equipment (steam, cooling water, compressed air, CIP) were not included.

In addition to the capital costs associated with the recovery process, operating costs were included in the model. These costs include chemicals (sulfuric acid for ion exchange regeneration, caustic for CIP), process consumables (membrane replacement costs, activated carbon, and IEX resin replacement costs), and utilities (steam, water, electricity, and wastewater). No additional labor burden was assumed in the model. Industrial-scale pricing for raw materials was used where available, and summary values are provided in **Table 3**.

**Table 3.**
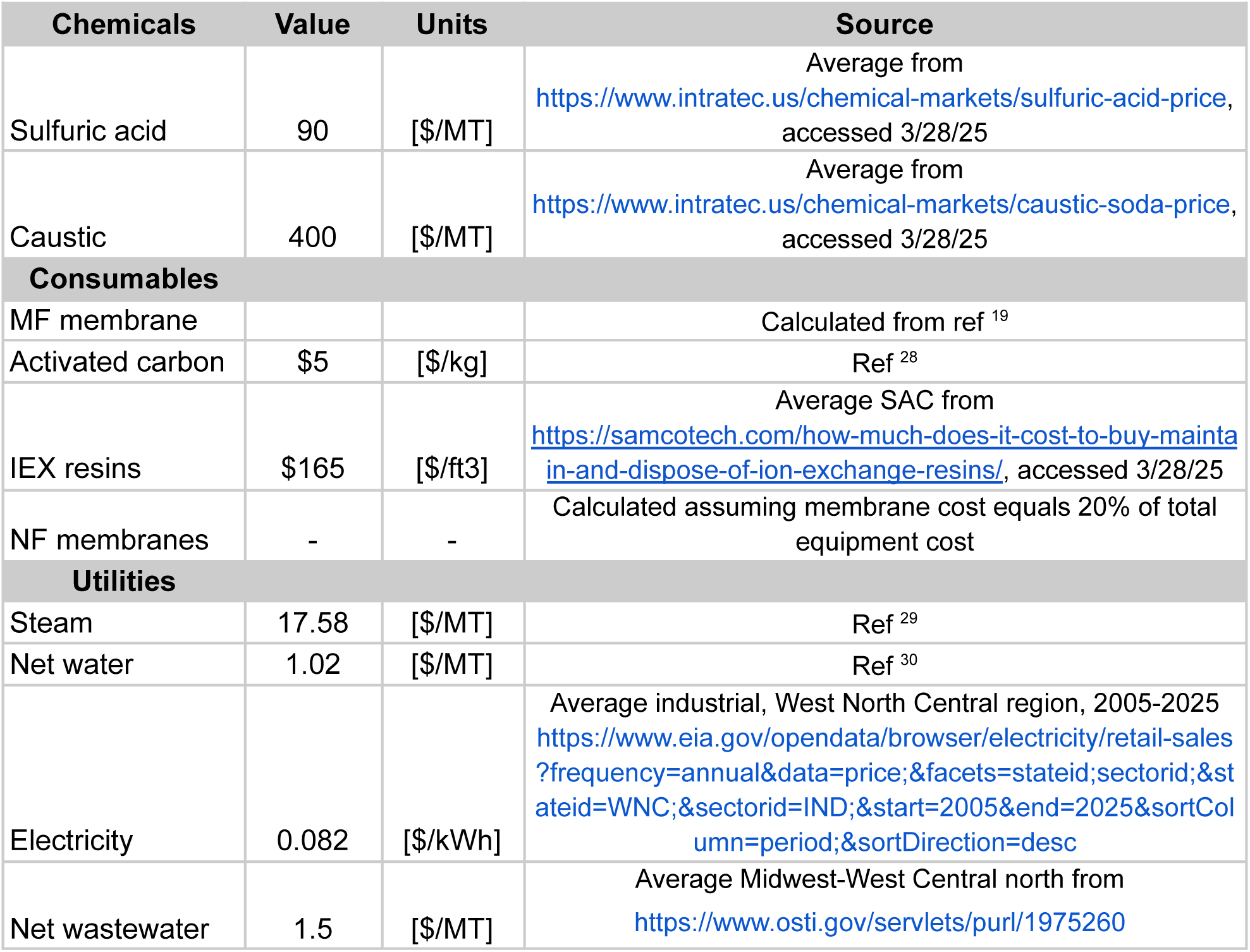
Operating costs used in the techno-economic model.

| Chemicals | Value | Units | Source |
| --- | --- | --- | --- |
| Sulfuric acid | 90 | [\$/MT] | Average from https://www.intratec.us/chemical-markets/sulfuric-acid-price, accessed 3/28/25 |
| Caustic | 400 | [\$/MT] | Average from https://www.intratec.us/chemical-markets/caustic-soda-price, accessed 3/28/25 |
| **Consumables** |  |  |  |
| MF membrane |  |  | Calculated from ref <sup>19</sup> |
| Activated carbon | \$5 | [\$/kg] | Ref <sup>28</sup> |
| IEX resins | \$165 | [\$/ft<sup>3</sup>] | Average SAC from https://samcotech.com/how-much-does-it-cost-to-buy-maintain-and-dispose-of-ion-exchange-resins/, accessed 3/28/25 |
| NF membranes | - | - | Calculated assuming membrane cost equals 20% of total equipment cost |
| **Utilities** |  |  |  |
| Steam | 17.58 | [\$/MT] | Ref <sup>29</sup> |
| Net water | 1.02 | [\$/MT] | Ref <sup>30</sup> |
| Electricity | 0.082 | [\$/kWh] | Average industrial, West North Central region, 2005-2025 https://www.eia.gov/opendata/browser/electricity/retail-sales?frequency=annual&data=price;&facets=stateid;sectorid;&stateid=WNC;&sectorid=IND;&start=2005&end=2025&sortColumn=period;&sortDirection=desc |
| Net wastewater | 1.5 | [\$/MT] | Average Midwest-West Central north from https://www.osti.gov/servlets/purl/1975260 |

## 7. Environmental impact analysis

An ex-ante attributional life cycle assessment (LCA) was performed to assess the environmental impact of the proposed lactic acid recovery process. A functional unit of 1 kg of 88% aqueous lactic acid was selected for the baseline analysis and the system boundary was set to include the lactic acid recovery process and all upstream raw material production and transport (**Figure 3**). Life cycle inventory (LCI) data for each unit operation were modeled using the Ecoinvent version 3.5 database in OpenLCA 1.11.0 software. The Midwest Reliability Organization (MRO) energy was selected to match energy production in the U.S. Midwest, where future cultivated meat facilities may be located ^31^. Process equipment for lactate recovery (e.g., evaporators, vessels, pumps) was not included, as the energy and resource impacts of their operation over a 15-year lifespan were assumed to dwarf the impacts from their materials, which can also be recycled at the end of life ^2^. The ReCiPe 2016 midpoint (H) and cumulative energy demand (CED) impact assessment methods were used to assess global warming potential (GWP), CED, and water use, and compared to values for lactic acid fermentation production in EcoInvent.

**Figure 3.**
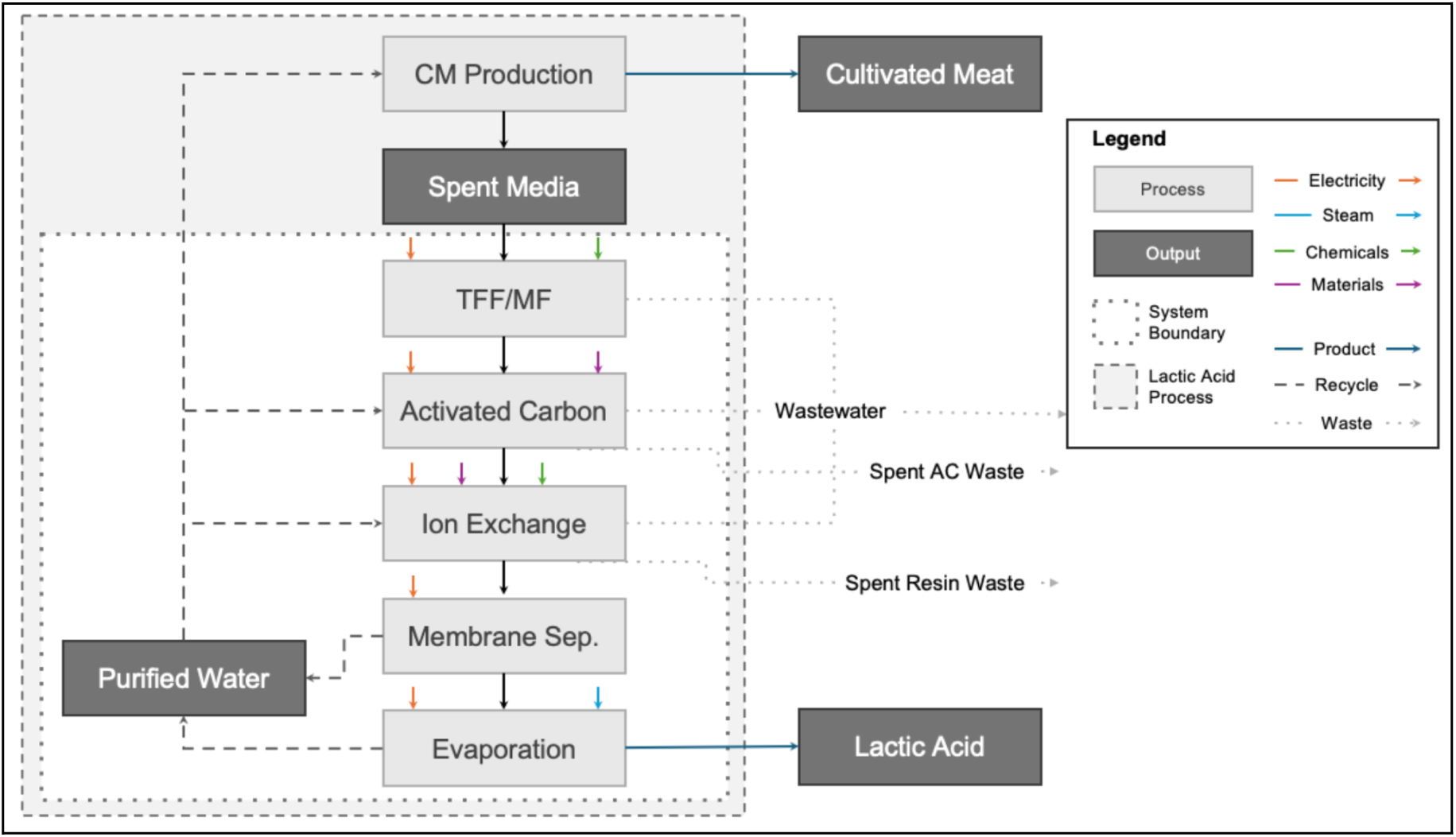
Overview of model and system boundaries for the LCA.

Values in Ecoinvent for lactic acid fermentation were on a 100% lactic acid basis and normalized to 88%. Information on allocation procedures is included in the **Supplementary Methods**. The full model and calculations can be found in the **Supplementary Spreadsheet**.

## 8. Sensitivity Analysis

Because of differences in spent media composition between cell lines and processes, opportunities for cultivated meat producers to modify lactate production, and expected improvements in total media use efficiency, a sensitivity analysis was carried out to explore the economic and environmental impacts of changes in lactic acid concentration and total media usage. A baseline concentration of 3 g of lactic acid per liter of spent media was used, with ranges of 1-5 g/L representing low and high scenarios, respectively. A baseline of 46.9 liters of spent media per kg of cultivated meat was used, with a ±25% difference applied to low and high scenarios. The economic allocation scenario was assumed for analysis of the effects on GWP and CED of cultivated meat production.

## Results

### Utilities for lactic acid recovery

In the baseline scenario with a lactic acid concentration of 3 g/L and a spent media flow rate of 58,597 L/hr, the recovery process results in a production rate of 127 kg of 88% aqueous polymer-grade lactic acid per hour, or 1,015 MTA. In the model facility, 0.1 kg of polymer-grade lactic acid is produced per kg of cultivated meat with an overall recovery of 64%.

The proposed process was estimated to consume approximately 2400 MT/yr of low-pressure steam for evaporation and to use approximately 1.4 million kWh of electricity for the operation of process equipment, driven primarily by NF pumps and cooling towers (**Figure 4**).

**Figure 4.**
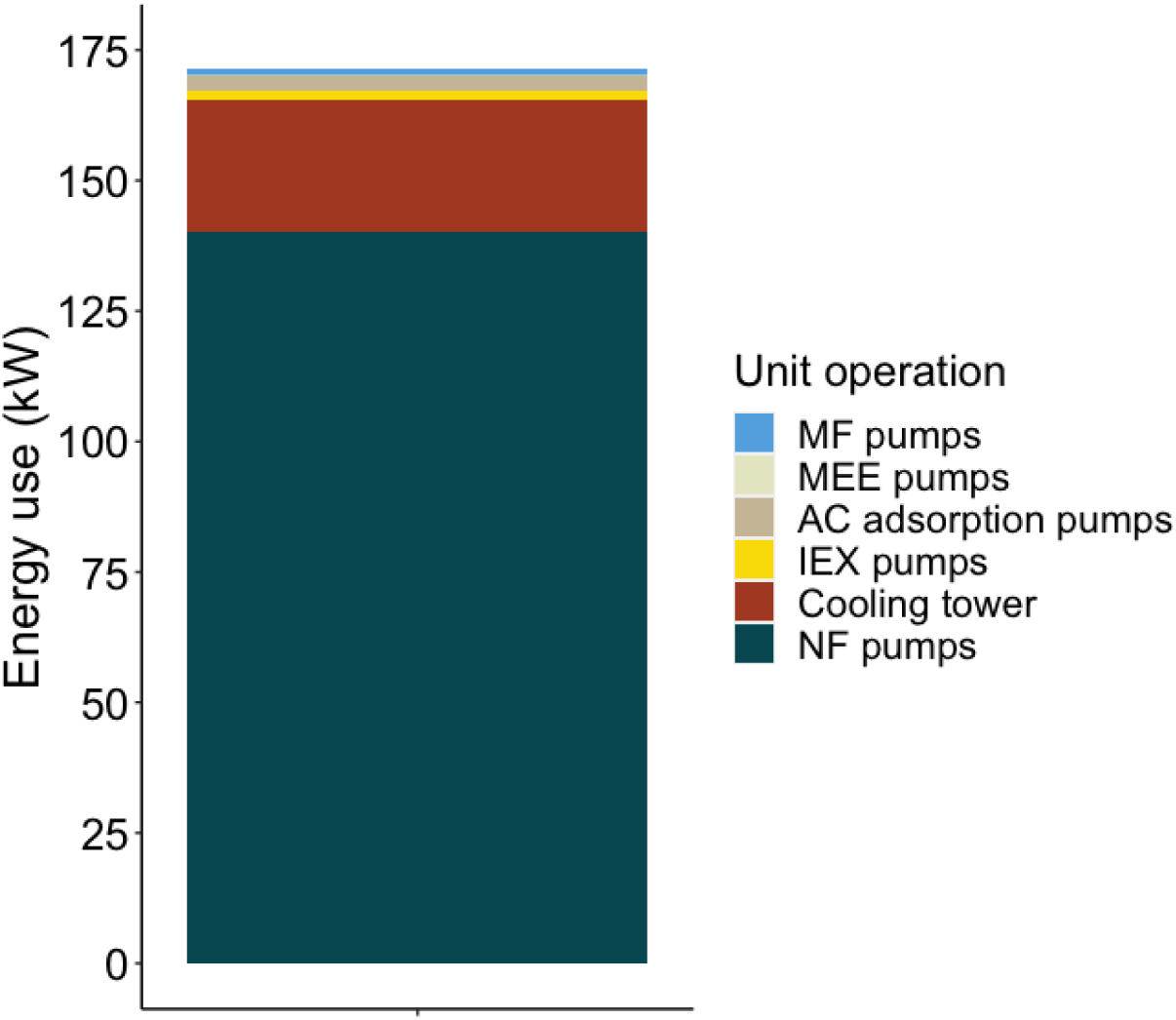
Annual energy consumption by unit operation in the recovery process. MF = microfiltration; MEE = multi-effect evaporator; AC = activated carbon; IEX = ion exchange; NF = nanofiltration.

Water use and wastewater production for the recovery process were calculated. The baseline process from Sinke et al was used to calculate net water usage and net wastewater requirements ^2^. While the NF and MEE lactic acid recovery steps produce a large amount of water (**Figure 1**, flows K and M, respectively) for reuse within the facility, the baseline process in Sinke et al assumed 75% water reuse, which is almost equal to the water reuse calculated in the recovery process model. As a result, the proposed process is estimated to be a small net consumer of water, equivalent to approximately 3 liters per minute (LPM), or 1,440 m^3^/yr (**Figure 5A**).

**Figure 5.**
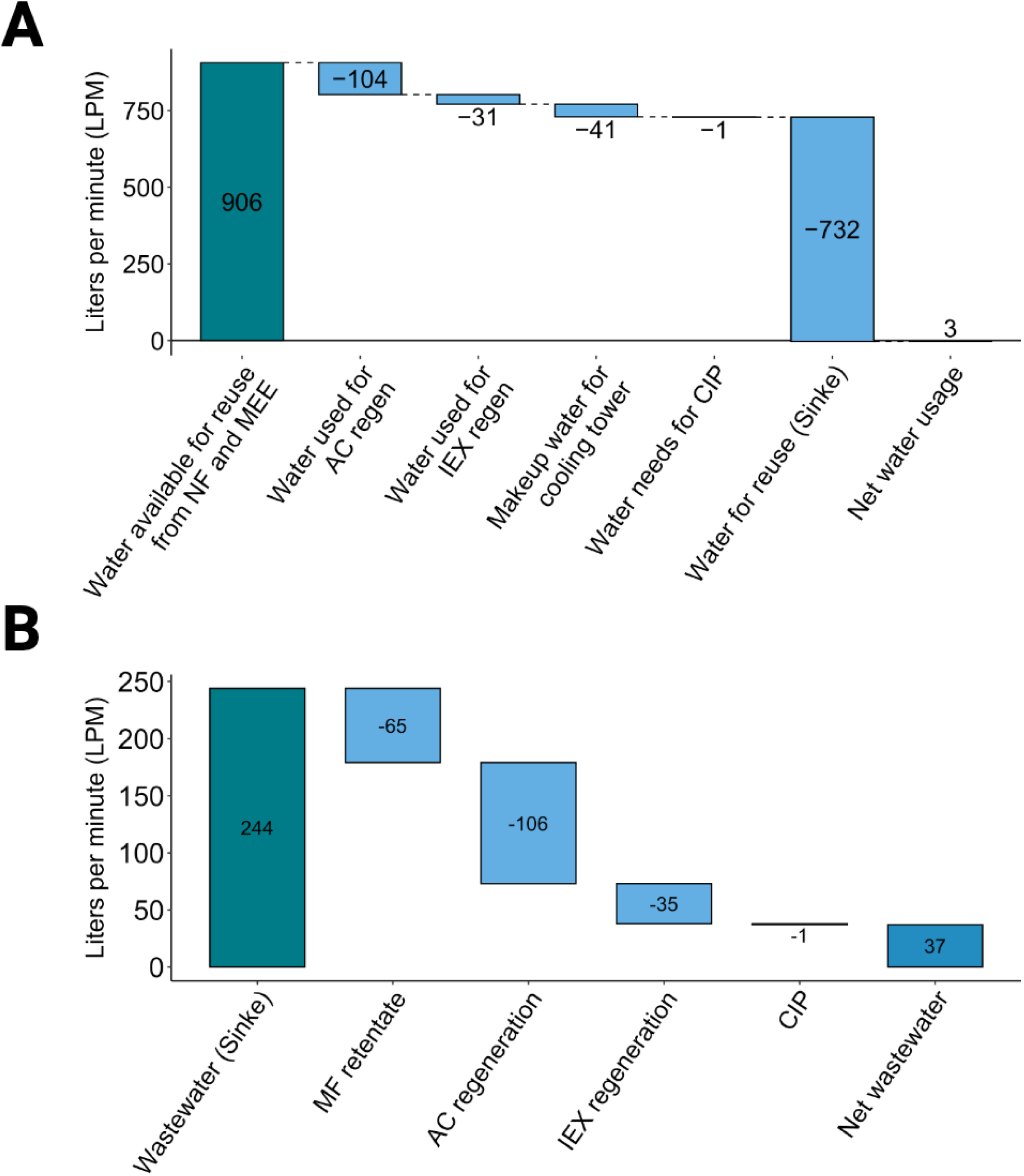
Water and wastewater use by unit operation. **(A)** Waterfall plot showing water use and reuse within the recovery process. **(B)** Waterfall plot showing wastewater production within the proposed process. MEE = multi-effect evaporator; AC = activated carbon; IEX = ion exchange; NF = nanofiltration; CIP = clean-in-place.

The reduction in required wastewater processing was also quantified, showing that the recovery process would reduce wastewater burden versus the baseline process by approximately 37 LPM, avoiding 17,760 m^3^/yr of wastewater treatment (**Figure 5B**). Credits for avoided wastewater were carried over to the operating costs and LCI associated with the integrated recovery process.

### Cost results

Capital costs were derived based on the sizing and installation of equipment for each unit operation. The largest capital costs were associated with the membrane separation of spent media and the removal of water ahead of evaporation (**Table 4**). With a 15-year flat depreciation schedule, capital charges with depreciation were calculated to be $0.31/kg of lactic acid (on an 88% lactic acid basis). The costs associated with membrane filtration may ultimately be avoided in future cultivated meat plants focused on the high recovery of proteins from spent media.

**Table 4.** Overview of capital costs for lactic acid recovery and purification. MEE = multi-effect evaporator; IEX = ion exchange; NF = nanofiltration; RO = reverse osmosis.

| Unit operation | Total cost (\$) | Percent of total capital cost |
| --- | --- | --- |
| Membrane filtration | \$2,157,464 | 46% |
| Activated carbon beds | \$293,017 | 6% |
| IEX | \$241,155 | 5% |
| NR/RO skid | \$1,018,552 | 22% |
| MEE | \$799,641 | 17% |
| Product storage | \$158,569 | 3% |
| **Total CAPEX** | **\$4,668,399** | **100%** |

Other lower-cost processes for media clarification, such as centrifugation, could also be explored for further cost reductions.

Operating costs associated with the recovery process were calculated, with costs being approximately equally distributed across the use of chemicals, consumables, and utilities (**Table 5**). Notably, the operational cost of water and wastewater resulted in a net savings, as only 3 LPM of water is used in the process, with 37 LPM of avoided wastewater providing a small credit.

**Table 5.** Overview of operating costs for lactic acid recovery and purification. MF = microfiltration; IEX = ion exchange; NF = nanofiltration.

| | Annual Cost<br>[\$] | Cost Contribution<br>[\$ /kg of 88% lactic acid] |
| --- | --- | --- |
| **Chemicals** |  |  |
| Sulfuric acid | \$145,191 | \$0.14 |
| Caustic | \$2,238 | \$0.00 |
| **Consumables** |  |  |
| MF membrane | \$26,781 | \$0.03 |
| Activated carbon | \$27,345 | \$0.03 |
| IEX resins | \$83,684 | \$0.08 |
| NF membranes | \$17,390 | \$0.02 |
| **Utilities** |  |  |
| Steam | \$42,000 | \$0.04 |
| Net water | \$25 | \$0.00 |
| Electricity | \$89,815 | \$0.09 |
| Net wastewater | -\$22,104 | -\$0.02 |
| **Total OPEX** | **\$412,365** | **\$0.41** |

At anticipated lactic acid concentrations of 3 g/L, the cost of recovery and purification to an aqueous solution of 88% lactic acid was calculated at approximately $0.71/kg. If sold at a competitive price of $1.41/kg, the annual production of 1,015 metric tons of lactic acid would result in a new revenue stream of $1.26MM/yr with an annual cost of production of $0.64MM/yr, resulting in $0.62MM in annual profit (**Table 6**). The installed capital has a calculated simple payback period of 7.5 years and profits from the sale of a lactic acid co-product could reduce cultivated meat COGS by approximately $0.06/kg.

**Table 6.** Financial metrics overview for the proposed lactic acid recovery process.

|  |  |
| --- | --- |
| **Annual revenue** | \$1.26MM |
| **Annual COGS** | \$0.64MM |
| **Annual profit** | \$0.62MM |

### Environmental impacts

The environmental impacts of the recovery process were analyzed using an ex-ante attributional LCA approach, focused on carbon footprint and energy use. The GWP and CED of recovering and purifying lactic acid were calculated to be 3.0 kg CO_2_eq and 44 MJ per kg of lactic acid, respectively. These values were approximately 1.0 kg CO_2_eq and 18 MJ less than the average values of lactic acid fermentation found in Ecoinvent (normalized on an 88% lactic acid basis), but not as low as some commercial processes ^32^.

Steam use during the evaporation stage accounted for over 50% of the impacts, followed by electricity use throughout the process and sulfuric acid needed for the regeneration of ion exchange columns (**Figure 6**). Other materials such as activated carbon, cleaning reagents, and wastewater had a negligible contribution.

**Figure 6.**
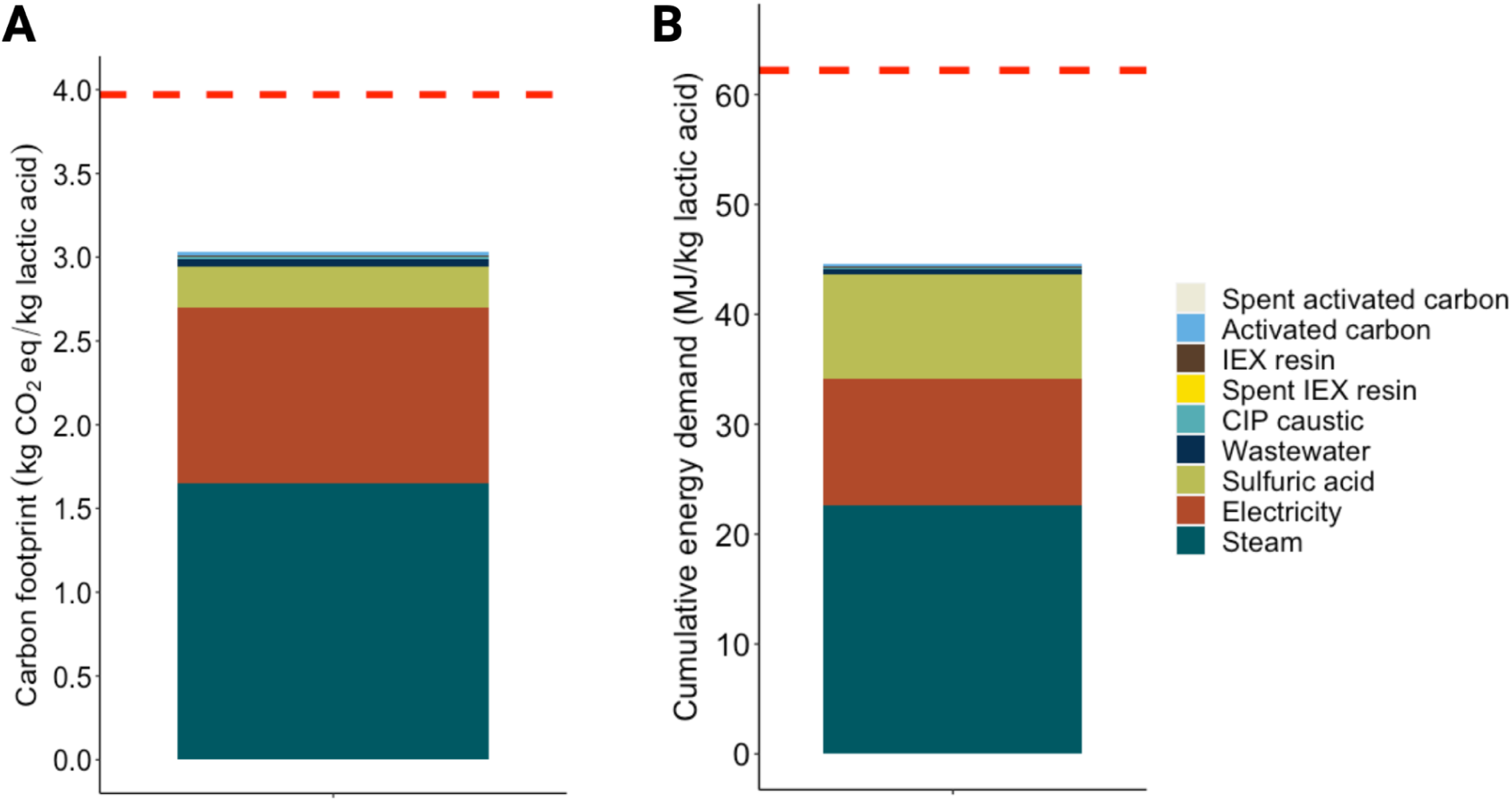
GWP and CED of the lactic acid recovery process. **(A)** Carbon footprint and **(B)** CED breakdown by unit operation. The red line represents the average GWP and CED of lactic acid production via fermentation.

Additional calculations were performed to contextualize the environmental impacts of recovered lactic acid as a co-product of cultivated meat production (**Table 8**). Incorporating the process into the baseline cultivated meat production scenario from Sinke *et al*, the cumulative GWP and CED of cultivated meat production with the integrated lactic acid recovery process become 14.5 kg CO_2_eq/kg and 283 MJ/kg of cultivated meat, respectively. Allocating impact based on economic value results in 14.4 kg CO_2_eq and 281 MJ assigned to cultivated meat and 0.1 kg CO_2_eq and 2 MJ assigned to lactic acid, respectively.

**Table 8.**
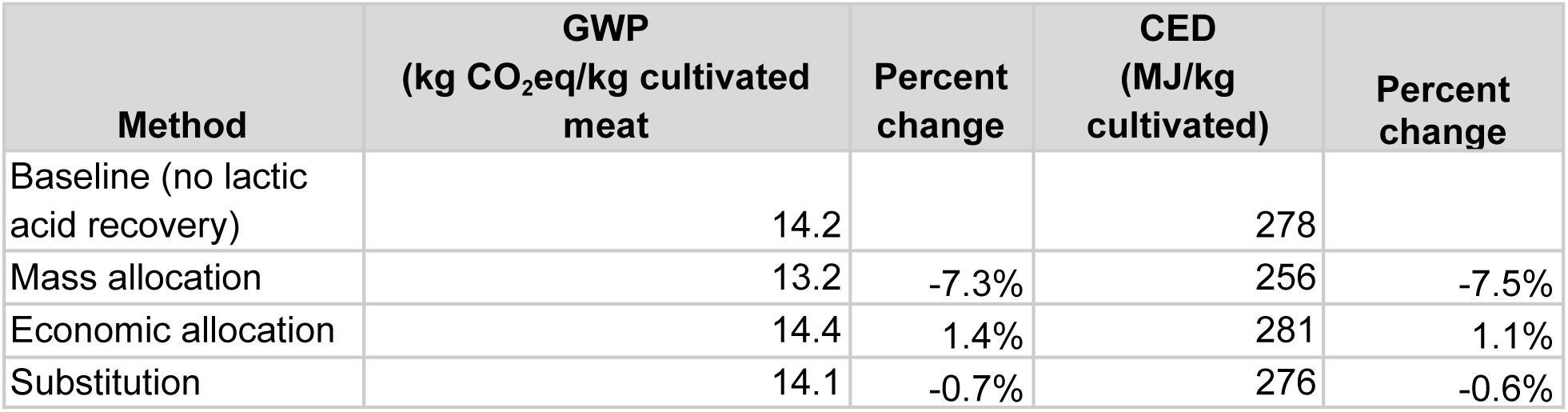
Summary of GWP and CED results of different allocation scenarios on cultivated meat production with integrated lactic acid recovery.

The impacts of lactic acid become extremely low because only 0.1 kg of lactic acid is produced per kg of cultivated meat, and the economic value of lactic acid is low compared to cultivated meat ($20/kg) in this analysis. Taken together, economic allocation from the perspective of a cultivated meat manufacturer employing lactic acid recovery in the facility modeled by Sinke *et al* results in a 0.2 kg CO_2_eq increase in carbon footprint and a 3.0 MJ increase in energy demand.

Allocation based on mass is an alternative way to contextualize the impacts. For completeness, the calculations of mass allocation were performed, resulting in a GWP of 13.2 kg CO_2_eq and a CED of 256 MJ per kg of cultivated meat. Thus, mass allocation from the perspective of a cultivated meat manufacturer employing lactic acid recovery in the facility modeled by Sinke *et al* is the most favorable, with a 1.0 kg CO_2_eq decrease in carbon footprint and a 22 MJ decrease in energy demand.

In the last method, the recovered lactic acid is assumed to displace virgin lactic acid produced via fermentation. This substitution has a favorable environmental impact, with a 1.0 kg CO_2_eq and 18 MJ reduction when compared directly to commercial lactic acid production. However, the yield of recovered lactic acid per kg of cultivated meat is low, resulting in the substitution scenario bringing a modest credit of-0.1 kg CO_2_eq and-1.8 MJ for each kg of cultivated meat produced, comparable to economic allocation.

Overall, depending on the allocation method, the estimated environmental impact of the recovery process had a modest-1.0 to +0.2 kg CO_2_ eq effect on the overall carbon footprint and a-22 to +3 MJ effect on cumulative energy demand per kg of cultivated meat production.

## Sensitivity analysis

Sensitivity analysis was performed to understand how results are affected when key parameters are changed. We analyzed scenarios where lactic acid concentration in spent media varied from 1 to 5 g/L and spent media production rates varied ±25% from baseline. For GWP and CED impacts on cultivated meat, the economic allocation scenario was assumed for baseline values.

The results show that the economics are highly sensitive to starting lactic acid concentration, with higher concentrations of 5 g/L affording significantly improved recovery costs at $0.43/kg of lactic acid, while lower concentrations of 1 g/L shift the COGS higher than the sales price of lactic acid, making recovery economically nonviable (**Figure 7**). Large shifts are also observed in the environmental impact of recovered lactic acid, where higher concentrations of lactic acid at 5 g/L result in a 42% decrease in GWP and CED, while lower concentrations of 1 g/L result in a 210% increase. However, the change in environmental impact on cultivated meat production remains modest, as the economic allocation method assigns most of the burden to the cultivated meat product (**Table 9**).

**Figure 7.**
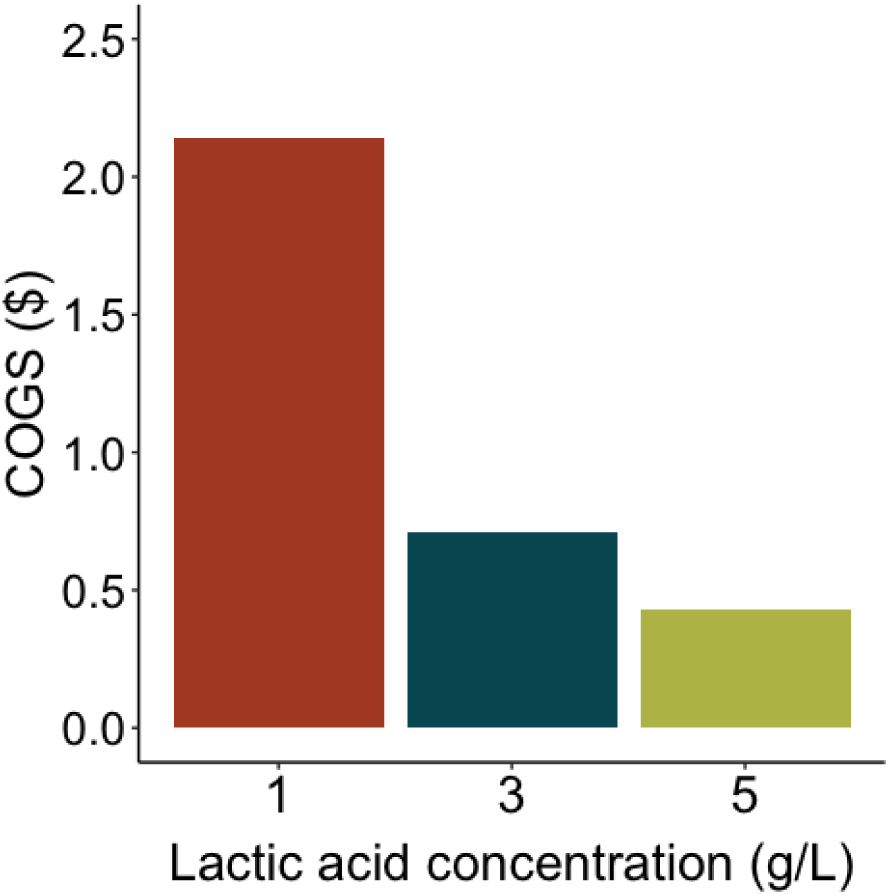
Sensitivity analysis of the COGS for lactic acid recovery when lactic acid concentration varies from 1 to 5 g/L.

**Table 9.** Overview of sensitivity analysis results for changes in starting lactic acid concentration. The top section shows the direct impact on recovered lactic acid, while the bottom section shows the impacts on cultivated meat, assuming economic allocation is applied. LA = lactic acid; CM = cultivated meat.

|  | Starting LA concentration (g/L) | GWP (kg CO<sub>2</sub>eq/kg LA) | GWP (% change) | CED (MJ/kg LA) | CED (% change) |
| --- | --- | --- | --- | --- | --- |
| GWP and CED impacts on recovered lactic acid | 1 | 9.3 | 210% | 137 | 210% |
|  | 3 (baseline) | 3.0 | - | 44 | - |
|  | 5 | 1.7 | -42% | 27 | -42% |
|  | Starting LA concentration (g/L) | GWP (kg CO<sub>2</sub>eq/kg CM) | GWP (% change) | CED (MJ/kg CM) | CED (% change) |
| GWP and CED impacts on cultivated meat, assuming economic allocation | 1 | 15.0 | 4% | 290 | 3% |
|  | 3 (baseline) | 14.4 | - | 281 | - |
|  | 5 | 14.3 | -1% | 279 | -1% |

Altering spent media production rates has minimal effects on COGS. While increasing the total processing rate may recognize small economies of scale in equipment costs, the COGS are weakly impacted, changing by 4% or less in either direction (**Figure S2**). Additionally, because the scale does not impact the normalized utility usage for lactic acid production, GWP and CED show no overall change with scale (**Table S1**).

## Discussion

Valorizing waste streams from industrial processes is a key theme to realizing a circular and sustainable future bioeconomy. Despite containing many nutrients and potentially valuable biomolecules, spent media from animal cell culture is currently treated as waste that is effectively flushed down the drain. In the biomanufacturing of antibody therapeutics alone, it has been estimated that at least 300 million liters of spent media are generated annually ^33^. The continued growth of cultivated meat, cellular therapeutics, and regenerative medicine industries will likely push total volumes into the billions of liters annually in the near future, adding stress to wastewater treatment facilities and limiting site selection of large-scale manufacturing plants only to municipalities that can handle the burden. A shift in perspective from spent media as waste to valuable sidestream or co-product is urgently needed.

With this in mind, we examined existing process technologies to conceptualize and design a five-step process to recover and purify the most abundant metabolite, lactic acid, from spent media. We found that recovery of lactic acid from spent media may provide a small improvement to both the economics and environmental impact of cultivated meat production. At an estimated net cost of $0.71 per kg of lactic acid, an integrated recovery process in a 10k MTA cultivated meat facility could yield an annual profit of $0.62MM, offsetting the cost of cultivated meat manufacturing by $0.06 per kg. The environmental impact of the recovery process was estimated to be 3.0 kg CO_2_eq and 44 MJ per kg of lactic acid, which was less than database values for commercial lactic acid fermentation processes. This may make purchasing recovered lactic acid attractive from the perspective of a downstream user, such as a food, cosmetics, or PLA manufacturer. However, the low yield of 0.1 kg of recovered lactic acid per kg of cultivated meat translates to minimal environmental savings for the cultivated meat manufacturer. Depending on how allocation is viewed, we found that lactic acid recovery could have a-1.0 to +0.2 kg CO_2_ eq effect on the overall carbon footprint and a-22 to +3 MJ effect on cumulative energy demand per kg of cultivated meat production.

This work aimed to create an objective evaluation of the economic and environmental impact of lactic acid recovery from spent media. While the proposed recovery process should provide high-purity lactic acid suitable for any downstream use, specific applications may have strict requirements necessitating further processing that is not accounted for here. Additionally, environmental data for lactic acid fermentation were normalized to an 88% lactic acid basis, which may be an oversimplification. However, these adjustments do not significantly impact the conclusions.

While the potential cost and environmental savings are modest, their relative impact should be placed in the appropriate context. As bioprocessing improves, cultivated meat manufacturing costs are likely to continually fall in the coming years, making even a few cents of improvement in COGS a meaningful change. Additionally, the baseline scenario for cultivated meat manufacturing in the Sinke study, which was used here, contained many conservative assumptions such as 75% of total energy use due to cooling demand for bioreactors ^2^. In practice, cooling may not be needed at all, resulting in significantly lower CED values that make relatively small savings from lactic acid recovery increasingly impactful. This relative increase in impact is also likely to occur as the share of renewable energy increases over time. For example, scenarios in the Sinke study using renewable energy had significantly lower GWP values of 2.5 - 4.0 kg CO_2_eq, and renewables would also be expected to lower the footprint of the recovery process, as a significant portion of its impacts are driven by electricity.

This study approached the separation problems inherent in lactic acid recovery using existing, mature technologies to provide a point of comparison and reference for future work on the valorization of waste streams in cultivated meat manufacturing. However, several exciting electrodialytic technologies are being developed that could further reduce recovery costs and environmental impacts ^34^. Additionally, custom resins using RNA show promise for highly selective separation and recovery of lactic acid^9^, while adsorbent technologies continue to advance for other potential coproducts such as ammonia^35^. While this study didn’t examine other technologies in detail, it is expected that improvements in membrane and adsorption technologies will continue to improve the sustainability and practicality of waste stream valorization processes.

### Future outlook for spent media recycling

Several other approaches are beginning to emerge for spent media recycling. The first approach involves using the separation and purification technologies described herein to remove metabolites such as lactate and ammonia before recycling the spent media back into the bioreactor where it is combined with fresh media to cultivate animal cells ^36^. Although these metabolites are problematic in cultivated meat production and typically considered waste, they also have mature commodity markets that could provide valorization opportunities, as demonstrated in this analysis. Another major benefit of this approach is the reuse of water, with the potential to carry over remaining nutrients and growth factors. This approach may be implemented in the near future for cultivated meat manufacturing, as the technology for practical application already exists and has been featured in industry patents ^21,37^.

The second approach involves using spent media as a feedstock or supplement for microbial production of food or pharmaceutical products. For example, spent media from CHO cells supplemented with 2% glycerol could support growth and recombinant protein production in *E. coli* equivalent to standard luria broth media ^33^. Similarly, spent media from chicken fibroblast cultures was shown to support growth factor production in *Lactococcus lactis* ^38^. Additionally, when used as a supplement, spent media from CHO cells outperformed typical fermentation broth for growing the fungi *Trametes versicolor* ^13^. These lab-scale proof-of-concept demonstrations suggest that spent media may be a cost-effective feedstock for the growing precision fermentation industry^39^, although further evidence at pilot scales and beyond will be needed.

The third approach involves using different microbes that consume lactate and ammonia to remediate spent media before recycling the treated media back into the animal cell culture. For example, the microalgae *Chlorella sorokiniana* was grown by consuming glucose and ammonia in spent media from quail myoblasts before the remediated media was recycled back into the quail cell culture ^40^. Variations of this approach involve the co-culture of microbes alongside animal cells. For example, a cyanobacterium strain was engineered to gain the function of lactate consumption and subsequently co-cultured in a transwell system to consume lactate and ammonia while returning pyruvate and amino acids to a rat cell culture. The conditioned media from the animal-cyanobacteria co-culture could then support the serum-free growth of mouse myoblast C2C12 cells ^41^. These studies collectively demonstrate that spent media from cultivated meat manufacturing — which would otherwise be viewed as waste that goes down the drain — can provide many necessary nutrients for growing microbial cultures of bacteria, microalgae, cyanobacteria, fungi, and potentially other organisms.

Future outlooks for a circular bioeconomy may entail certain components of spent media being targeted for valorization or remediation, while other bulk media nutrients support microbial growth for the purposes of food ingredient production. The resulting microbial biomass can be subsequently hydrolyzed into a nutrient supply to be fed back into the animal cell culture system (**Figure 9**). Although more research will be needed to determine the most suitable microbes, hydrolysis protocols, and bioprocess designs that implement media recycling in this fashion, laboratory demonstrations using hydrolyzed microalgae and bacterial lysates have already provided some promise ^42^ ^43^.

**Figure 9.**
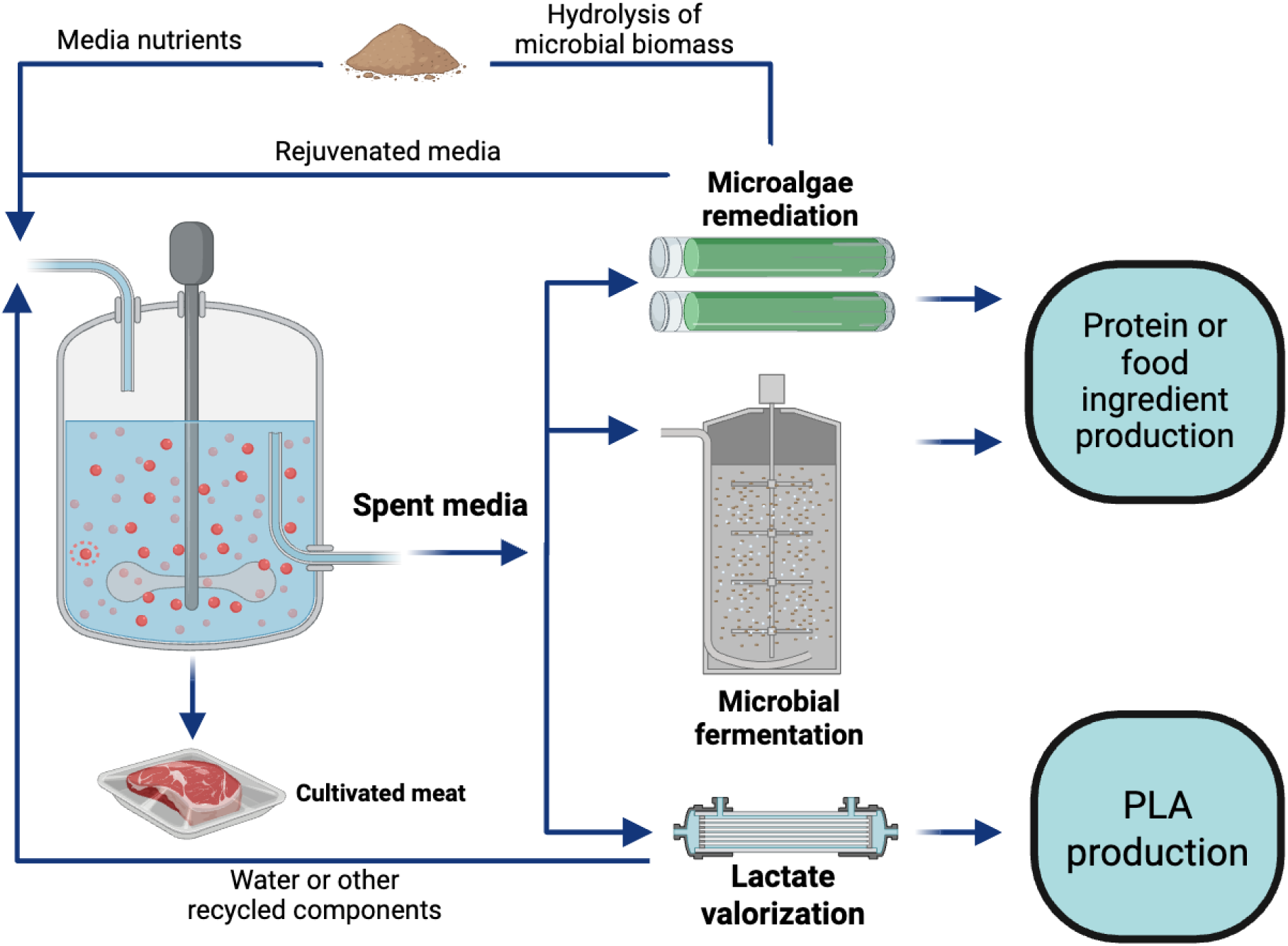
Future outlook for a circular bioeconomy within the broader cellular agriculture industries.

Cultivated meat manufacturers may consider co-localizing their production facilities alongside bioplastics or microbial production that occurs under the same roof or next door, saving on transport costs of liquid spent media without the need to dehydrate for shipping over long distances. As more emphasis on the bioeconomy develops in the coming years, cooperation amongst a broad range of stakeholders, including industry, policymakers, governments, and NGOs, will be needed to realize these approaches at scale.

## Conclusion

As the cultivated meat industry continues to expand, new strategies are needed to reduce the cost and environmental impact of production. This study presents a combined techno-economic and environmental impact assessment for the recovery of lactic acid from spent media in cultivated meat manufacturing. By conceptualizing and modeling a five-step process based on commercially available separation and recovery technologies, we demonstrate that lactic acid valorization from spent media is not only technically feasible but could also deliver modest economic and environmental benefits.

While our model results are strongly influenced by lactic acid concentrations in the spent media, baseline values from other studies suggest that the proposed process strategy may provide economic recovery of lactic acid for a wide range of cell types and processes. Our baseline model shows a path to recovery of lactic acid at $0.71/kg, which could generate $0.06 in value per kg of cultivated meat, creating a new revenue stream and potentially offsetting a portion of the production costs. Life cycle assessment further shows that the recovery of lactic acid can improve GWP and CED by up to ∼7%, depending on how the allocation is contextualized.

Beyond cultivated meat production, the findings have broader implications for other sectors of animal cell culture. As cell culture processes continue to scale, valorizing abundant metabolites such as lactic acid and ammonia presents actionable steps toward circular biomanufacturing.

While this study relied on established separation technologies, future work should explore novel approaches to recovery and how they may fit into integrated facilities. As other industries continue to face cost constraints and explore opportunities to valorize waste streams, the conceptual approach described here can provide a framework for considering the impacts of different coproducts and recovery schemes. Taken together, this work suggests that integration of lactic acid recovery into next-generation cultivated meat facilities should be considered in more detail, laying the groundwork for broader adoption of waste stream valorization across the industry.

## Author Contributions

**Joshua Wimble**: Conceptualization; formal analysis; methodology; writing—original draft; writing—review and editing. **Reina Ashizawa**: Formal analysis; methodology; writing—original draft. **Elliot Swartz**: Conceptualization; methodology; writing—original draft; writing—review and editing; project administration; supervision.

## Acknowledgments

This work was funded by the generous donors who support the Good Food Institute, a nonprofit science-driven think tank helping to build a more sustainable, secure, and just food system.

We thank Amin Nikkah for guidance in environmental impact calculations and Adam Leman for critical review and feedback on the manuscript.

Figure 9 and Figure S1 were created in https://BioRender.com.

