## Supplemental Figures and Methods for "Analysis of the economic viability and environmental impacts of a conceptual process for the recovery of lactic acid from spent media in cultivated meat production"

**Supplementary Methods**

### Additional considerations for recovery and purification of lactic acid from spent media

One consideration for lactate recovery and purification is the potential for biomolecules such as alcohols and organic acids to interfere with the process or result in quality challenges for PLA production. For example, alcohols can react with lactic acid via esterification reactions producing compounds such as ethyl lactate. However, animal cells generally lack the metabolic pathways to produce alcohols, limiting this as a concern for lactic acid recovery from spent media.

Other organic acids such as citric acid, acetic acid, pyruvic acid, or acidic amino acids can be produced by cells and could be found in spent media at detectable concentrations, although likely at least 100-1000x lower than lactic acid. Having similar chemical properties as lactic acid, these molecules may compete for binding sites on ion exchange (IEX) resins, alter the pH, or contribute to membrane fouling. Additionally, large molecular weight entities such as large proteins, cell debris, and exosomes could impact purification processes by contributing to filter fouling and end product quality. Accordingly, the five-step process was conceptualized to be able to work with these limitations in mind.

### Allocation rationale and methods

Different scenarios were used to evaluate the potential impacts of lactic acid recovery in the context of the overall cultivated meat process. These scenarios included cultivated meat production within the system boundary, with a functional unit of 1 kg of cultivated meat. In the first scenario, impacts were economically allocated because the lactic acid is intended to be sold as a revenue-generating co-product of cultivated meat production. The price of cultivated meat was set at $20/kg based on what may be attainable in a 10k MTA facility within a five-year timeframe (Pasitka et al. 2024), while 100% lactic acid was set at $1.60/kg, which was the average of lactic acid prices cited in various reports (Biddy, Scarlata, and Kinchin 2016; Zacharof and Lovitt 2013). The price of 88% lactic acid was calculated from this value to be $1.41. Economic allocation was chosen based on the significant revenue disparity between cultivated meat and lactic acid, aligning with EN 15804 guidelines that state a >25% revenue difference justifies the use of economic allocation (The British Standards Institution 2012). Calculations for mass allocation were also performed for comparison.

In the second scenario, system expansion was applied by assuming recovered lactic acid displaces the production of virgin lactic acid from fermentation, enabling evaluation of the environmental benefit of displacing its production. This approach was chosen because the recovery process yields 88% aqueous polymer-grade lactic acid that is intended to match the purity, performance, and functionality of commercial fermentation-derived lactic acid, allowing for a 1:1 substitution. Average inventory data for the substituted lactic acid was used rather than marginal data because the study follows an attributional framework and the recovered lactic acid represents a small fraction of total commercial production. Given the multi-billion-dollar scale of the lactic acid market, quantities of lactic acid recovered from cultivated meat production are not expected to disrupt market dynamics in the near future.

Because the lactic acid recovery process generates water for reuse through nanofiltration and distillation, the net water use was considered in the LCI. After calculating the total water recovered, a 75% water reuse rate, based on Sinke et al., was applied to calculate the net impact on water consumption and wastewater generation in the 10k MTA plant. The LCI of water use includes GWP and CED values for the attributional and system expansion analyses. Similar calculations were performed for wastewater generation.

**Supplemental figures and tables**

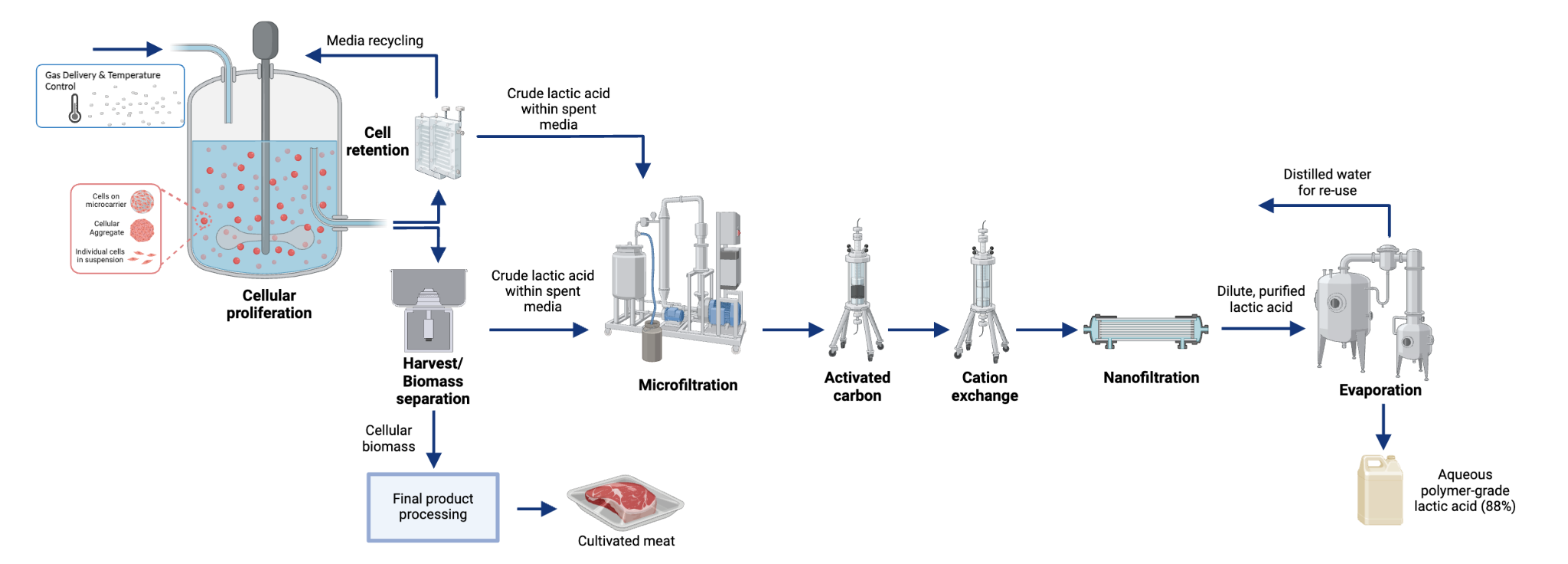

**Figure S1.** Integrated process for lactic acid recovery from spent media during cultivated meat production, resulting in an aqueous polymer-grade lactic acid (88%) co-product. Spent media can be harvested from perfusion or batch processes and passed through the five-step recovery process using existing membrane separation and adsorption technologies. A multi-stage evaporation process is used to concentrate the lactic acid, with recovered water available for reuse in the facility. Recovered lactic acid can then be bottled and sold to customers, resulting in an additional revenue stream for cultivated meat manufacturers.

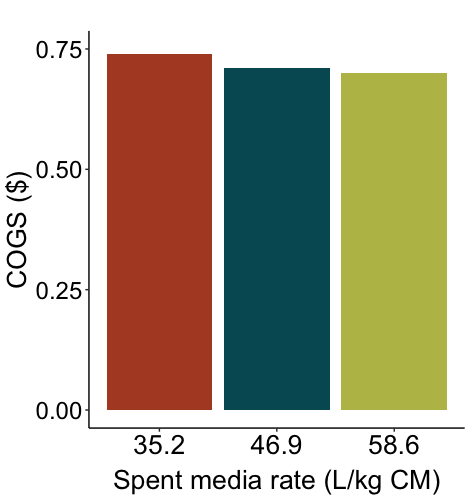

**Figure S2.** Impact of the COGS for lactic acid recovery when spent media production rate varies from 35.2 to 58.6 L/kg cultivated meat. CM = cultivated meat.

|  | **Spent media rate  (L/kg CM)** | **GWP  (kg CO2eq)** | **GWP  (% change)** | **CED  (MJ)** | **CED  (% change)** |
| --- | --- | --- | --- | --- | --- |
| GWP and CED impacts on cultivated meat assuming economic allocation | 35.2 | 14.4 | 0% | 281 | 0% |
|  | 46.9 (baseline) | 14.4 | - | 281 | - |
|  | 58.6 | 14.4 | 0% | 281 | 0% |

**Table S1.** Overview of sensitivity analysis results for GWP and CED impacts on cultivated meat, assuming economic allocation, based on changes in spent media rate.
